# The tempo and mode of angiosperm mitochondrial genome divergence inferred from intraspecific variation in *Arabidopsis thaliana*

**DOI:** 10.1101/769653

**Authors:** Zhiqiang Wu, Gus Waneka, Daniel B. Sloan

**Author notes:** Authors contributed equally (Zhiqiang Wu, Gus Waneka). Corresponding author: Daniel B. Sloan, 1878 Campus Delivery, Fort Collins, CO 80523.

## Abstract

The mechanisms of sequence divergence in angiosperm mitochondrial genomes have long been enigmatic. In particular, it is difficult to reconcile the rapid divergence of intergenic regions that can make non-coding sequences almost unrecognizable even among close relatives with the unusually high levels of sequence conservation found in genic regions. It has been hypothesized that different mutation/repair mechanisms act on genic and intergenic sequences or alternatively that mutational input is relatively constant but that selection has strikingly different effects on these respective regions. To test these alternative possibilities, we analyzed mtDNA divergence within *Arabidopsis thaliana*, including variants from the 1001 Genomes Project and changes accrued in published mutation accumulation (MA) lines. We found that base-substitution frequencies are relatively similar for intergenic regions and synonymous sites in coding regions, whereas indel and nonsynonymous substitutions rates are greatly depressed in coding regions, supporting a conventional model in which mutation/repair mechanisms are consistent throughout the genome but differentially filtered by selection. Most types of sequence and structural changes were undetectable in 10-generation MA lines, but we found significant shifts in relative copy number across mtDNA regions for lines grown under stressed vs. benign conditions. We confirmed quantitative variation in copy number across the *A. thaliana* mitogenome using both whole-genome sequencing and droplet digital PCR, further undermining the classic but oversimplified model of a circular angiosperm mtDNA structure. Our results suggest that copy number variation is one of the most rapidly evolving features in angiosperm mtDNA, even outpacing rearrangements in these notoriously structurally diverse genomes.

## INTRODUCTION

The evolution of angiosperm mitochondrial genomes (mitogenomes) is a study in contrasts. On one hand, they exhibit exceptionally low nucleotide substitution rates, including at synonymous sites even though such sites are likely subject to relatively low levels of functional constraint (WOLFE *et al*. 1987; DROUIN *et al*. 2008). These low levels of sequence divergence are generally assumed to reflect unusually slow point mutation rates, especially when compared to high mitochondrial mutation rates in many other eukaryotic lineages (BROWN *et al*. 1979; SLOAN *et al*. 2017). However, direct measures of plant mitochondrial mutation rates are generally lacking, and the mechanisms that maintain such low levels of nucleotide substitutions are not known.

On the other hand, angiosperm mitogenomes are remarkably diverse at a structural level (MOWER *et al*. 2012b; GUALBERTO and NEWTON 2017). They are large and variable in size and subject to extensive rearrangements via recombination-mediated mechanisms, which may be accelerated under conditions of plant stress (ARRIETA-MONTIEL and MACKENZIE 2011). Although they typically map as circular structures, their actual physical form appears to be far more complex and variable (BENDICH 1993; SLOAN 2013; KOZIK *et al*. 2019).

Comparisons among angiosperm mitochondrial genomes often find that large fractions of intergenic sequence are unalignable between species and seemingly unique to individual lineages (KUBO and NEWTON 2008). In the most extreme cases, only about half of intergenic sequence content may be shared even between two different mitochondrial haplotypes from the same species (SLOAN *et al*. 2012). There are likely at least two mechanisms responsible for this phenomenon. First, angiosperm mitogenomes are frequent recipients of large quantities of horizontally transferred DNA from the plastid genome, nucleus, and other sources (ELLIS 1982; GOREMYKIN *et al*. 2012; RICE *et al*. 2013). As such, many intergenic sequences are recently acquired and truly lack homologous sequences in mitogenomes of other angiosperms. It is unlikely, however, that horizontal transfer can provide a full explanation because a lot of intergenic content cannot be traced to any potential donor source. A second possible mechanism is that rates of sequence and structural evolution are so fast in the intergenic regions of angiosperm mitogenomes that homologous sequences can become essentially unrecognizable even among closely related species. But this latter explanation presents a paradox when juxtaposed with the observation that genic regions in plant mitogenomes can exhibit some of the slowest known rates of nucleotide substitutions.

Christensen (2013; 2014) has proposed alternative models to explain the striking contrast in evolutionary rates between genic and intergenic regions in angiosperm mitogenomes, which are based either on differences in mutational input or differences in selection between these two types of regions. Under the mutational-input model, the contrasting rates of divergence would reflect systematic differences between genic and intergenic sequences with respect to DNA polymerase errors during replication, exposure to DNA damage, and/or the efficacy of DNA repair processes. It was hypothesized that transcription-coupled repair (HANAWALT and SPIVAK 2008) could have such an effect in altering mutation rates in expressed vs. non-expressed regions in angiosperm mitogenomes (CHRISTENSEN 2013), but subsequent analysis of substitution rates in transcribed non-coding regions did not find support for this hypothesis (CHRISTENSEN 2014). Nevertheless, the possibility of systematic differences in mutational input among regions within plant mitochondrial genomes remains largely untested, and it has been observed that some species can exhibit substantial rate variation even from one gene to the next for reasons that remain unclear (ZHU *et al*. 2014; WARREN *et al*. 2016).

An alternative and perhaps more conventional model is that mutational input is relatively consistent across the genome but that genic vs. intergenic regions are subject to very different selection pressures. For example, structural and sequence variation introduced by error prone repair pathways may be filtered out in gene regions but largely neutral and tolerated in non-coding regions (CHRISTENSEN 2014). This may be especially true for any repair mechanisms that lead to structural rearrangements or indels that would truncate protein-coding genes. One prediction from this model is that rates of single-nucleotide substitutions in intergenic regions should largely match those at relatively neutral sites in protein-coding sequences (e.g., synonymous sites). However, this prediction has been difficult to test because finding sets of genomes that have enough divergence in coding regions to estimate substitution rates and still retain enough similarity in intergenic structure and content to align these non-coding regions is a challenge.

In this sense, variation at an intraspecific scale may be informative, as comparisons between patterns of recent and long-term evolutionary change can be powerful in separating effects of mutation and selection (NIELSEN 2005). A previous pairwise comparison between two different *Arabidopsis thaliana* accessions was used to measure mitochondrial sequence divergence, but this analysis only identified a single synonymous nucleotide substitution in protein-coding genes and thus could offer little precision in quantifying the frequency of single nucleotide polymorphisms (SNPs) in different functional sequence categories (CHRISTENSEN 2014). The study was further complicated by the large number of sequencing errors that were later identified in the early *A. thaliana* mitogenome reference sequences (SLOAN *et al*. 2018).

Here, we take advantage of the ever-growing amount of genomic resources in *A. thaliana*, including the sequencing of complete genomes from the 1001 Genomes Project (ALONSO-BLANCO *et al*. 2016) and from mutation accumulation (MA) lines in this species (JIANG *et al*. 2014), to generate more robust polymorphism datasets for investigating the mechanisms of mitogenome divergence. Our goal is to distinguish among alternative explanations for the contrasting rates of genic vs. intergenic sequence evolution and identify the genomic changes that accrue most rapidly during angiosperm mitogenome evolution.

## MATERIALS AND METHODS

### Identification of intraspecific mitogenome variation from the *Arabidopsis* 1001 Genomes Project

To analyze standing mitochondrial polymorphisms within *A. thaliana*, raw Illumina reads from the 1001 Genomes Project (which actually contains 1135 sequenced individuals; ALONSO-BLANCO *et al*. 2016) were downloaded from the NCBI Sequence Read Archive (SRA) under the project accession SRP056687 using the fastq-dump tool in the NCBI SRA Toolkit v2.9.6. For larger datasets, only the first 20 million read pairs were downloaded. Illumina adapter sequences were trimmed with Cutadapt v2.1 (MARTIN 2011), applying a q20 quality cutoff, a 15% error rate for matching adapter sequences, and a minimum trimmed read length of 50 bp. As such, 88 of the 1135 sequenced individuals were excluded entirely from the analysis because their original read lengths were shorter than 50 bp. Trimmed reads were mapped to the *A. thaliana* Col-0 GenBank RefSeq accessions for the mitochondrial (NC_037304.1) and plastid genomes (NC_000932.1) using Bowtie v2.3.5 (LANGMEAD and SALZBERG 2012). By competitively mapping sequence reads against both organelle genomes, we avoided erroneously mapping plastid-derived reads to related regions in the mitogenome resulting from historical plastid-to-mitochondrial DNA transfers (i.e., *mtpts*; ELLIS 1982; SLOAN and WU 2014). The resulting alignment files were sorted with SAMtools v1.9 (LI *et al*. 2009), and variants were called using the HaplotypeCaller tool in GATK v4.1.0.0 (MCKENNA *et al*. 2010) with ploidy level set to 1 after removing duplicate reads with the GATK MarkDuplicates tool. Coverage depth at each position in the mitogenome was calculated with the SAMtools depth function. The resulting variant sets were filtered to require a minimum site-specific coverage depth of 50. Variants were also excluded if their coverage was less than half or more than three times the median genome-wide coverage. These thresholds were applied to avoid erroneously identifying variants based on low-frequency sequences such as nuclear insertions (i.e., *numts*; STUPAR *et al*. 2001; HAZKANI-COVO *et al*. 2010) or based on mis-mapping to repeats within the genome.

To distinguish between ancestral and derived alleles that are segregating within *A. thaliana*, we aligned the *A. thaliana* reference genome against the *Brassica napus* mitogenome (NC_008285.1), using NCBI BLASTN v2.2.30+, applying a minimum alignment length of 400 bp and a minimum nucleotide identity of 90%. The *B. napus* allele for all alignable *A. thaliana* SNP positions was extracted from the BLAST output with a custom BioPerl script (STAJICH *et al*. 2002), which is available via GitHub (https://github.com/dbsloan/polymorphism_athal_mtdna). An alternative approach to distinguish between ancestral and derived alleles is based on the fact that derived alleles are typically at low frequency. As such, even when it is not possible to polarize a variant with an outgroup because it is found in an unalignable region, reasonable predictions of ancestral vs. derived state can still be based on current allele frequencies. Therefore, we calculated allele frequencies at each variable site to identify the minor allele, using all samples within the 1001 Genomes Project that met our coverage requirements for variant calling (see above).

Positions within the *A. thaliana* reference mitogenome were partitioned into functional categories (protein-coding, rRNA, tRNA, introns, pseudogenes, and intergenic) based on the RefSeq annotation (NC_037304.1). PAML v4.9a was used to approximate the total number of synonymous and nonsynonymous ‘sites’ within protein-coding sequence (accounting for the partial degeneracy at some positions owing to two- and three-member codon families).

### Analysis of mitogenome divergence in *Arabidopsis* mutation accumulation lines

To analyze short-term divergence in *A. thaliana* mitogenomes, we obtained raw Illumina reads from the MA lines generated by Jiang et al. (2014) from NCBI SRA (SRP045804). MA lines involve bottlenecking each generation through single-seed descent to limit selection on organismal fitness and obtain a relatively unfiltered view of *de novo* mutation accumulation (HALLIGAN and KEIGHTLEY 2009). This dataset consisted of a total of six MA lines, each propagated for 10 generations. Three lines were propagated under benign growing conditions, while the other three were subjected to salt stress each generation, except in the final generation in which all lines were grown under the same benign conditions. Three biological replicates from each of the six lines were sequenced in the original study (JIANG *et al*. 2014).

To test for *de novo* nucleotide substitutions and indels in the mitogenomes of these MA lines, we applied the same variant calling pipeline as described above for the 1001 Genomes samples. The only modification was that we set the ploidy level to 10 so that we could potentially detect any novel variants that were heteroplasmic at a frequency of ∼10% or greater. There are many causes that can lead to erroneous identification of *de novo* mitochondrial variants, including mapping artefacts, *numts*, and heteroplasmies inherited from the original parent. To avoid such errors, we focused on variants that were unique to one or more replicates from a single MA line. For all such variants predicted by our pipeline, we manually inspected read alignments using IGV (ROBINSON *et al*. 2017) to determine whether they were detectable in samples from other MA lines.

We analyzed copy number variation across the *A. thaliana* mitogenome by normalizing site-specific data for depth of sequence coverage as counts per million mapped read (CPMM) values and averaging them into non-overlapping windows of 500 bp. To avoid any effects of cross-mapping from plastid-derived reads, which are highly abundant in total-cellular DNA samples, we excluded any windows that overlapped with previously identified *mtpts* (SLOAN and WU 2014). We also excluded the first and last windows because of potential bias in mapping at the edges where the circular mitogenome map was arbitrarily cut into a linear sequence. To try to account for coverage bias introduced during the sequencing process because of differences in local nucleotide composition (AIRD *et al*. 2011; VAN DIJK *et al*. 2014), we fit these data to a linear model that included GC content and a count of homopolymers of greater than 7 bp in length as independent variables to predict CPMM in each window. This model was implemented in R v3.6.0 using the lm function. The subsequent analyses of copy number variation described below were performed with both the raw CPMM values and the residuals from this model.

To test for associations in coverage values between adjacent windows across the mitogenome, we performed a Wald–Wolfowitz runs test, using the runs.test function in the R randtests package. To test for significant divergence in coverage values among the MA lines, we fit a model with treatment (salt-stressed vs. control) as a fixed effect and MA line as a nested random effect. This test was implemented in R with the lmer function and the lme4 and lmerTest R packages. We controlled for multiple comparisons by applying a false discovery rate (FDR) correction (BENJAMINI and HOCHBERG 1995).

We also examined the frequency of alternative genome conformations associated with recombination between small repeats by first mapping Illumina reads to the *A. thaliana* Col-0 reference mitogenome with BWA v0.7.12, using the mem command and the -U 0 option. We then used a custom Perl script (https://github.com/dbsloan/polymorphism_athal_mtdna) to parse the resulting alignment file. For each pair of repeats in the mitogenome, this script calculated the number of read pairs that mapped in a concordant fashion spanning a repeat as well as the number of read pairs that mapped discordantly but in locations that were consistent with a recombination event between the pair of repeats. This analysis was performed on all repeat pairs between 100 and 500 bp in length with a minimum of 80% nucleotide sequence identity. We then tested whether the frequency of recombinant conformations for each repeat pair differed significantly among MA lines by once again fitting a model with treatment as a fixed effect and MA line as a nested random effect (see coverage analysis described above).

### Mitochondrial DNA purification and Illumina sequencing

Three full-sib families from our *A. thaliana* Col-0 lab stock were grown in a growth chamber under short-day conditions (10 h of light at 100 µmole m^-2^ s^-1^) at 23 °C. For each family, 30-40g of rosette tissue was harvested from plants after 6-7 weeks of growth. To reduce starch content, plants were kept in the dark for two days prior to collecting leaf tissue, and then the harvested tissue was stored overnight in the dark at a 4 °C. All subsequent tissue-processing and DNA-extraction steps were carried out in a 4 °C cold room or refrigerated centrifuge unless stated otherwise.

Leaf tissue was disrupted in high salt isolation buffer (1.25 M NaCl, 50 mM Tris-HCl pH 8.0, 5 mM EDTA, 0.5% polyvinylpyrrolidone, 0.2% bovine serum albumin, 15 mM β-mercaptoethanol), using 10 ml of buffer per g of tissue. Disruption was performed with a standard kitchen blender and a series of five bursts of ∼10 s each with ∼10 s of settling time between each burst, followed by filtration through four layers of cheesecloth and one layer of Miracloth. Filtrates were then centrifuged at 150 rcf for 15 min. The resulting supernatant was transferred to new bottles and centrifuged at 1500 rcf for 20 min. The supernatant was then again transferred to new bottles and centrifuged at 15,000 rcf for 20 min. After discarding the resulting supernatant, the mitochondrial pellets, were gently but thoroughly resuspended in 3 ml of DNase buffer (0.35 M sorbitol, 50 mM Tris-HCl pH 8.0, 15 mM MgCl_2_) with a paintbrush. Then 7 ml of DNase solution (DNase I dissolved in DNase buffer at a concentration of 1 mg/ml) was added to each resuspended pellet. The samples were incubated on ice for 1 h with occasional gentle swirling to digest contaminating plastid and nuclear DNA. Three volumes of wash buffer (0.35 M sorbitol, 50 mM Tris-HCl pH 8.0, 25 mM EDTA) was added to each sample followed by centrifugation at 12,000 rcf for 20 min. The resulting pellets were washed two more times by resuspending in 20 ml wash buffer and centrifuging at 12,000 rcf for 20 min. The final washed pellet was resuspended in 1 ml wash buffer. One-twentieth volume of a 20 mg/ml proteinase K solution was added and incubated at room temperature for 30 min. Mitochondria were lysed by adding one-fifth volume of lysis buffer (5% N-lauryl sarcosine Na salt; 50 mM Tris-HCl pH 8.0, 25 mM EDTA) followed by gentle mixing by inversion for 10 min at room temperature. One volume of phenol:chloroform:isoamyl alcohol (25:24:1) was added followed by vortexing for 5 s and centrifugation at 12,000 rcf for 10 min. The resulting aqueous phase was transferred to a new tube and incubated with 4 µl of a 10 mg/ml RNase A solution. The samples were then treated with two rounds of cleanup with phenol:chloroform:isoamyl alcohol as described above followed by precipitation with one volume of ice-cold isopropanol and incubation for at least 20 min at −20 °C. Precipitated DNA was pelleted by centrifugation at 12,000 rcf for 10 min and washed twice with 500 µl of ice-cold 70% ethanol. The final DNA pellet was air dried and dissolved in TE buffer (10 mM Tris-HCl pH 8.0, 1 mM EDTA).

Sequencing libraries were produced for each of the three resulting mtDNA samples, using the NEBNext Ultra II FS DNA Library Prep Kit. We used 50 ng of input DNA, with a 15 min fragmentation step, and 5 cycles of PCR amplification. The resulting libraries had an average insert size of approximately 245 bp and were sequenced on a NovaSeq 6000 platform (2×150 bp), producing between 14.1M and 15.4M read pairs per library. The reads were used for coverage-depth analysis by mapping to the *A. thaliana* reference mitogenome as described above for the MA-line dataset.

### ddPCR copy number analysis

To confirm variation in copy number that was inferred from deep sequencing data across the mitogenome, we performed droplet digital PCR (ddPCR). Primers were designed to target six regions with high sequencing coverage and six regions with low coverage (Table S1). Analysis, was performed on the same three purified mtDNA samples described above and one sample of total-cellular DNA extracted from the same *A. thaliana* Col-0 lab line, using a modified CTAB and phenol:chloroform protocol (DOYLE and DOYLE 1987). The template quantity for each reaction was either 2 pg of mtDNA or 400 pg of total-cellular DNA, with two technical replicates for each reaction. All ddPCR amplifications were set up in 20-μL volumes with Bio-Rad QX200 ddPCR EvaGreen Supermix and a 2 μM concentration of each primer before mixing into an oil emulsion with a Bio-Rad QX200 Droplet Generator. Amplification was performed on a Bio-Rad C1000 Touch Thermal Cycler with an initial 5 min incubation at 95 °C and 40 cycles of 30 s at 95 °C and 1 min at 60 °C, followed by signal stabilization via 5 min at 4 °C and 5 min at 95 °C. The resulting droplets were read on a Bio-Rad QX200 Droplet Reader. Copy numbers for each PCR target were calculated based on a Poisson distribution using the Bio-Rad QuantaSoft package. To assess significant difference in copy-number between the sets of primers from high- and low-coverage regions of the mitogenome, one-tailed t-tests were performed for each of the four DNA samples, using the means for each pair of technical replicates as input.

### Data Availability

All newly generated and previously published sequence data are available via NCBI SRA. Newly generated Illumina data were deposited under accession PRJNA546277. Custom scripts used in data analysis are available via GitHub (https://github.com/dbsloan/polymorphism_athal_mtdna). Data pertaining to identified sequence variants and copy-number variation are provided in supplementary Figures S1-S4 and Tables S1-S4 submitted via https://gsajournals.figshare.com.

## RESULTS

### Intraspecific mitochondrial sequence variation in the *Arabidopsis thaliana* 1001 Genomes Project

Using whole-genome resequencing data from the 1001 Genomes Project, we identified a total of 1105 mitochondrial SNPs that are variable across *A. thaliana* accessions, including three sites at which three different alleles were detected (Table S2). For a subset of 319 of these sites, we could infer the ancestral state by aligning the nucleotide position to the outgroup *Brassica napus*. We could also infer the polarity of changes for the entirety of the dataset by assuming that minority allele represented the derived state. This allele-frequency method produced the same call for 87% of the 319 *Brassica*-polarized SNPs, suggesting that it had substantial predictive value. Both of these approaches revealed a mutation spectrum that is heavily biased towards increasing AT content. Substitutions that increased AT content were 7-fold more common than those that decreased it based on the *Brassica*-polarized dataset and 5-fold more common in the full dataset based on allele frequency (Table S2). The spectrum did not exhibit the large overrepresentation of transitions that is found in mtDNA of some eukaryotes (YANG and YODER 1999), with an overall transition:transversion ratio of 422:686 that was only modestly above the null expectation of 1:2 (Table S2). However, AT→TA and GC→CG transversions were rare, representing only 7% and 10% of all transversions, respectively (Table S2). This mutation spectrum is generally consistent with observations from a published pairwise comparison between the *A. thaliana* Col-0 and C24 ecotypes (CHRISTENSEN 2013). The extreme AT bias is also consistent with a previous analysis of inserted plastid sequences (*mtpts*) as relatively neutral markers in angiosperm mtDNA (SLOAN and WU 2014). Although that study found that angiosperm mitogenomes generally had weak AT bias, it identified *A. thaliana* as an outlier with a much stronger bias than most species. Therefore, the inferred mitochondrial mutation spectrum from *A. thaliana* may not be broadly representative of angiosperms with respect to AT bias.

By comparing the distribution of SNPs across different functional classes within the mitogenome, we found that the presence of base-substitutions is 2.9-fold lower in protein-coding and RNA genes than in intergenic regions (Table 1). However, if only synonymous SNPs in protein-coding genes are considered, the SNP abundance is much more similar but remains slightly lower in genes (0.0027 per synonymous site) than in intergenic regions (0.0034 per site). The average minor allele frequency was also slightly lower for synonymous SNPs (0.016) than for SNPs in intergenic regions (0.026).

**Table 1.** Variant statistics for 1001 Genomes dataset. SNPs: single nucleotide polymorphisms; MAF: minor allele frequency.

| Sequence Type | Sites | SNPs | SNPs per Site | SNP MAF | Indels | Indels per Site | Indel MAF |
| --- | --- | --- | --- | --- | --- | --- | --- |
| Protein Coding | 31264 | 41 | 0.0013 | 0.0206 | 0 | 0.0000 | NA |
| Nonsynonymous | 24323 | 22 | 0.0009 | 0.0244 | 0 | 0.0000 | NA |
| Synonymous | 6941 | 19 | 0.0027 | 0.0163 | 0 | 0.0000 | NA |
| rRNA | 5222 | 3 | 0.0006 | 0.0010 | 0 | 0.0000 | NA |
| tRNA | 1689 | 0 | 0.0000 | NA | 0 | 0.0000 | NA |
| Pseudogene | 1256 | 5 | 0.0040 | 0.0025 | 0 | 0.0000 | NA |
| Intron | 35335 | 72 | 0.0020 | 0.0218 | 18 | 0.0005 | 0.0116 |
| Intergenic | 293042 | 987 | 0.0034 | 0.0263 | 172 | 0.0006 | 0.0239 |
| **Total** | **367808** | **1108** | **0.0030** | **0.0256** | **190** | **0.0005** | **0.0006** |

In contrast to the relatively similar SNP levels between synonymous sites and intergenic regions, there was a radical difference in the distribution of indels across functional classes in the *A. thaliana* mitogenome. A total of 190 polymorphic indels were identified in the 1001 Genomes dataset, and every one of them was located in either an intergenic region or an intron (Table 1). Overall, within gene sequences, we found a large reduction of variants that are expected to be disruptive of gene function (i.e., nonsynonymous substitutions and indels) but limited evidence of reduced abundance of changes that are likely to be relatively neutral (i.e., synonymous substitutions).

### Shifts in mitochondrial copy-number variation across mutation accumulation lines

By analyzing mitochondrial reads from published whole-genome resequencing data of *A. thaliana* MA lines (JIANG *et al*. 2014), we found that most potential mitogenome changes were undetectable over a timescale of 10 generations, regardless of whether the lines had been propagated under salt-stressed or benign conditions. We did not detect any SNPs or indels that reached homoplasmy in individual lines. Our pipeline identified a total of 11 low-frequency variants (seven SNPs, two indels, and two multinucleotide variants with multiple changes clustered at nearby sites) that were unique to a single MA line and thus candidates for *de novo* mutations. However, manual inspection of read alignments found evidence of these same variants at low frequencies in other lines, indicating that they were unlikely to be true *de novo* mutations. Therefore, we did not find any convincing evidence of novel substitutions or small indels present in the heteroplasmic state. Angiosperm mitogenomes are known to undergo frequent, homogenizing recombination between large repeat sequences and lower frequency recombination between short repeat sequences (<500 bp), which can lead to shifts in the relative frequency of alternative structures (SMALL *et al*. 1987; LONSDALE *et al*. 1988; ARRIETA-MONTIEL *et al*. 2009; GUALBERTO and NEWTON 2017). To test for such structural changes, we quantified the frequency of recombinant conformations using read-pairs spanning short repeat sequences. Although we identified minor variation in frequencies of alternative conformation across sequenced lines (Table S3), none of these showed consistent patterns of divergence for either treatment or line effects at an FDR of 0.05.

In mapping MA line reads to the *A. thaliana* reference mitogenome, we observed variation in coverage across the length of the genome, which was broadly similar in the six different MA lines (Figure 1). Because Illumina DNA sequencing (and the PCR-based techniques it relies on) can be biased against sequences with extreme GC or AT richness or with low-complexity features like homopolymers (AIRD *et al*. 2011; VAN DIJK *et al*. 2014), it is possible that the observed coverage variation was an artefact of amplification/sequencing bias. To investigate this possibility, we fit a model to predict sequencing depth based on GC content and presence of homopolymers. This effort was only able to explain a low percentage of the variance in sequencing depth across the mitogenome (*R*^2^ < 0.3 for all datasets), and the general pattern of copy number of variation was retained after controlling for this effect (Figure S1), suggesting that bias associated with simple nucleotide-composition features was not the primary cause of the observed variation. For subsequent analyses of coverage depths, we also used the residuals from these models to account for sequencing bias related to nucleotide composition.

**Figure 1.**
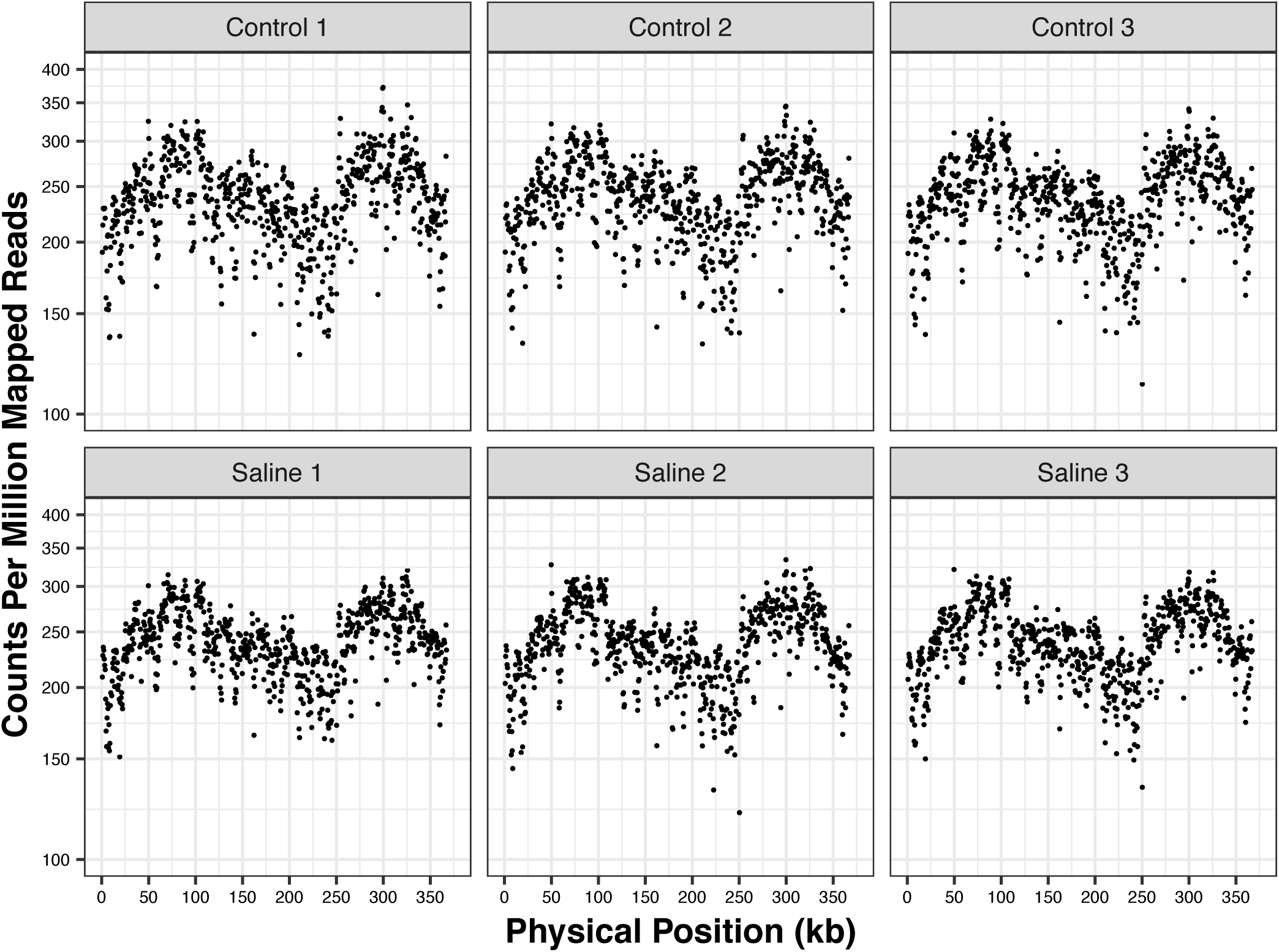
Sequencing coverage variation across mitogenome of *Arabidopsis thaliana* mutation accumulation lines. Each panel represents an average of three biological replicates.

To assess whether there were any significant shifts in copy-number variation during propagation of MA lines, we scanned the length of the genome in 500-bp windows to test for effects at the level of treatment (salt-stressed vs. control) and individual MA lines. We found that many of the 713 windows in the mitogenome showed small but significant differences between treatments after an FDR correction for multiple comparisons (35 windows when using raw CPMM values and 14 when using residuals from a nucleotide composition model; Figures 2 and S2; Table S4). None of the windows were significant for an MA-line effect after the same FDR correction, where line was tested as a nested effect within treatment (Table S4). Adjacent regions tended to show coverage differences in the same direction relative to the genome-wide median (Wald–Wolfowitz runs test; only 201 observed cases in which adjacent windows were on opposite sides of the median compared to a null expectation of 356; *P* << 0.001). Therefore, we found evidence that MA lines shifted in consistent ways with respect to region-specific copy number when subjected to stressed vs. benign growing conditions over 10 generations. Although the effect sizes were modest (up to a 20.5% shift in coverage in stressed vs. control samples), they could still be detected with a relatively small sample size because of the consistent patterns across replicate lines.

**Figure 2.**
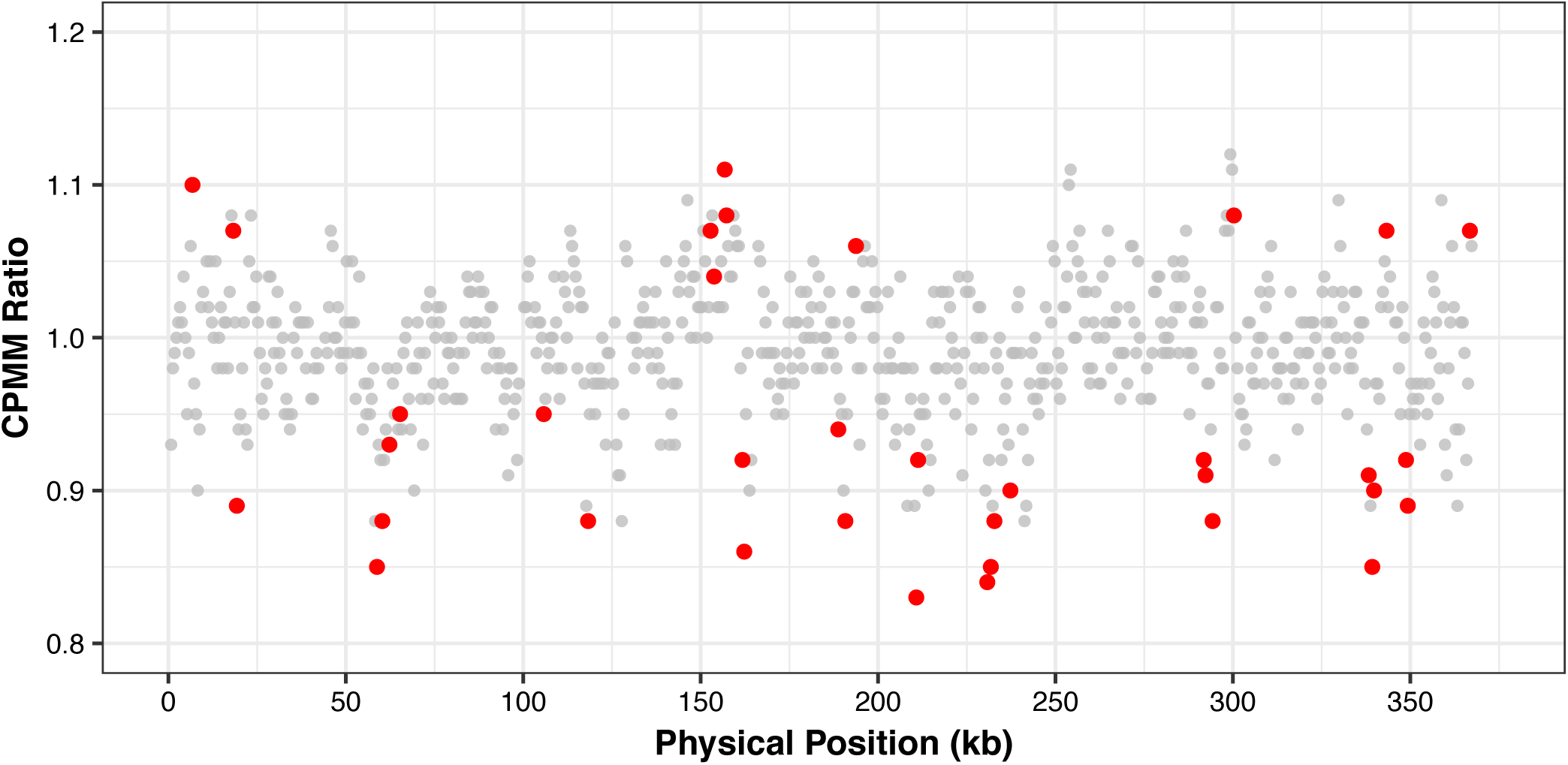
Divergence in region-specific mitogenome copy number in salt-stressed vs. control mutation accumulation lines. Values are expressed as a ratio of the averages for all salt-stressed and all control lines. Windows that deviate significantly from a ratio of 1 after false-discovery-rate correction are highlighted in red. CPMM: counts per million mapped reads.

### Sequencing and ddPCR analysis of purified *Arabidopsis thaliana* mtDNA

To further test for evidence of copy number variation within the *A. thaliana* mitogenome, we purified mtDNA from replicate families of our own lab line of the Col-0 ecotype. Illumina sequencing of these samples resulted in ∼60% of reads mapping to the *A. thaliana* reference mitogenome, demonstrating substantial enrichment for mtDNA. As found with the MA lines, this analysis revealed a heterogeneous pattern of coverage across the mitogenome, which was generally consistent among the three replicates (Figures 3 and S3). Once again, we found that adjacent regions tended to show coverage variation in the same direction (Wald–Wolfowitz runs test; only 120 observed cases in which adjacent windows were on opposite sides of the median compared to a null expectation of 356; *P* << 0.001). However, comparing between our samples and the MA lines found only a modest correlation in copy number variation (*r* < 0.25; Figure S4).

**Figure 3.**
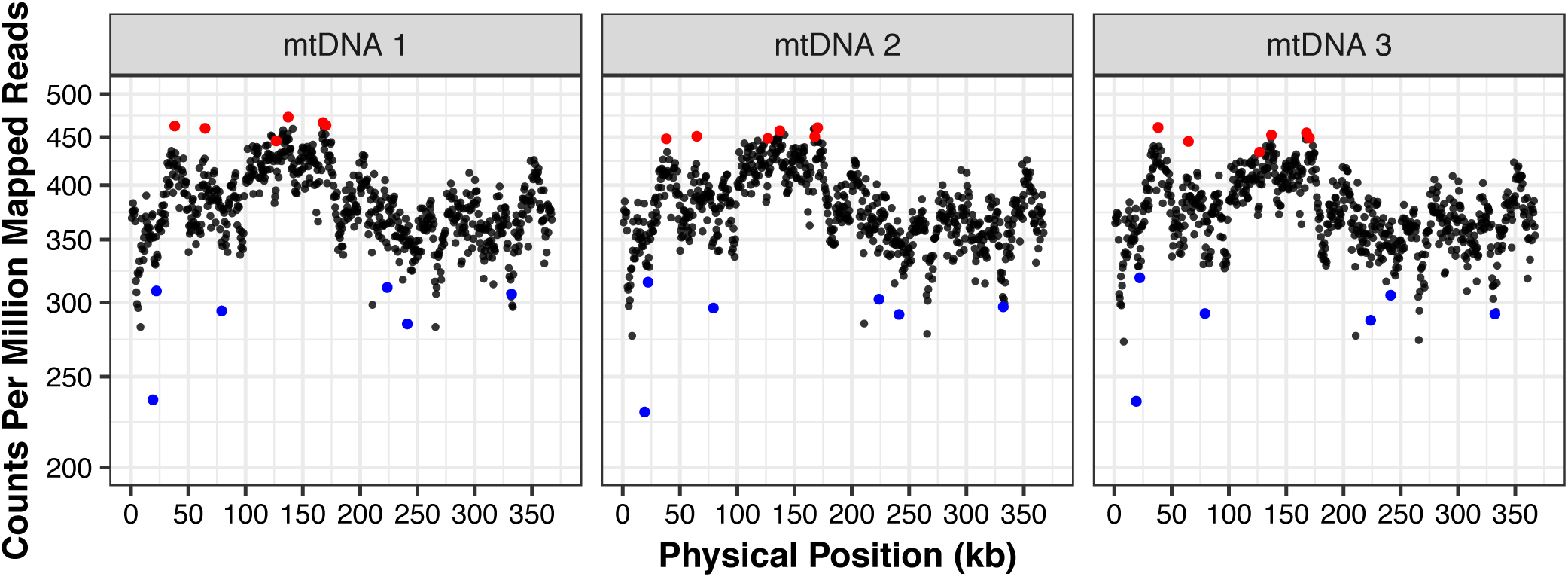
Sequencing coverage variation across the mitogenome for three purified mtDNA samples from *Arabidopsis thaliana*. The windows chosen for development of ddPCR markers are shown in red and blue dots (high- and low-coverage regions, respectively).

To confirm that the observed heterogeneity in coverage was a product of true variation in copy number rather than an artefact of sequencing bias, we performed ddPCR with two sets of six markers that were selected for either high-coverage or low-coverage regions based on sequencing data (Figure 3). Unlike sequencing and traditional qPCR, this method is generally insensitive to variation in PCR efficiency or amplification bias because it is based on endpoint PCR (40 cycles) within each ‘micro-reactor’ droplet. We found significant differences in copy number between the sets of high- and low-coverage markers for both the purified mtDNA samples that were used in sequencing and a total-cellular DNA extraction (*P* < 0.001 for each of the three purified mtDNA samples and *P* = 0.011 for the total-cellular DNA sample; Figure 4). In all cases, the average difference in copy number between these sets was somewhat smaller (between 17.1% and 20.3% for the purified mtDNA samples and 10.7% for the total-cellular sample) than from sequence estimates (mean of 36.3%), which may reflect some regression to the mean because the high- and low-copy markers were chosen only based on being in the extreme tails of the sequencing-coverage distribution rather than for an *a priori* reason.

**Figure 4.**
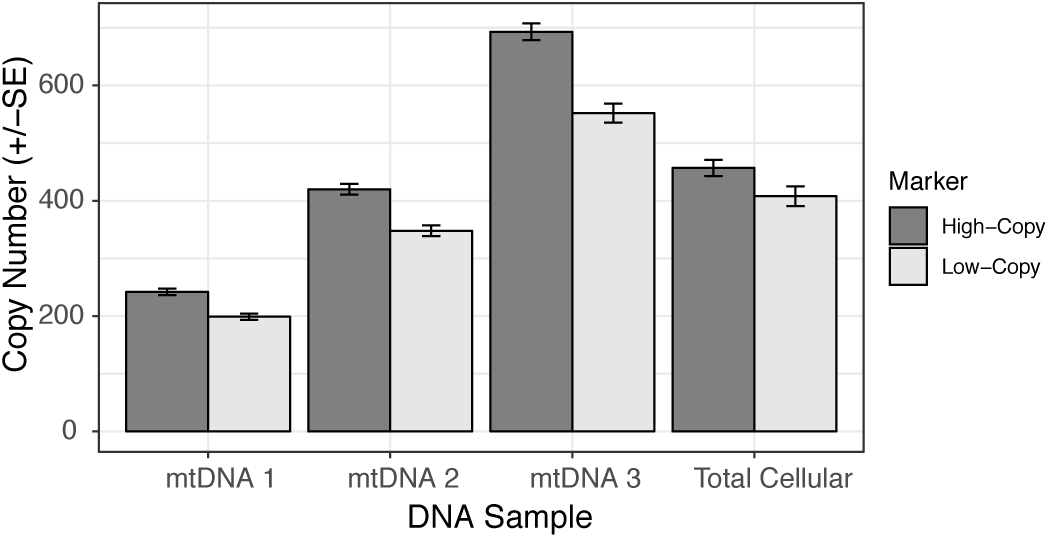
ddPCR comparison of copy number for mitogenome regions identified as either high-copy or low-copy by sequencing analysis. Copy numbers are expressed as per µl of ddPCR reaction volume. Input for the mtDNA samples was diluted 200-fold relative to the total-cellular sample.

## DISCUSSION

### Contrasting rates of evolution in genic and intergenic regions of angiosperm mitogenomes

Our analysis confirmed dramatic differences in rates of mitogenome structural evolution between genic and intergenic regions at an intraspecific level within *A. thaliana*, mirroring the extensive observations of this phenomenon based on divergence between angiosperm species (KUBO and NEWTON 2008). By dramatically expanding the number of sampled accessions with the aid of the 1001 Genomes dataset (ALONSO-BLANCO *et al*. 2016), we were also able to make quantitative comparisons between nucleotide substitution rates in these regions, which was previously difficult because of the limited number of substitutions in an earlier comparison between two *A. thaliana* accessions (CHRISTENSEN 2013). The similar levels of nucleotide substitutions between intergenic regions and synonymous sites in protein-coding genes (Table 1) suggests that mutational input in different functional regions is comparable. As such, the most likely explanation for the divergent evolutionary rates in genic and intergenic regions is a conventional model, under which selection has varying effects in filtering mutations in different region throughout the genome (CHRISTENSEN 2014).

Despite the rough similarity between nucleotide substitution rates at synonymous sites and in intergenic regions, we still found that the synonymous rate was slightly lower (Table 1). There are multiple possible explanations for this gap. First, it is possible synonymous substitution rates are suppressed because these sites still experience a (larger) degree of purifying selection. For example, even if they do not change amino acid sequences, synonymous substitutions can disrupt the translation efficiency, secondary structure, or binding motifs of mRNAs (CHAMARY *et al*. 2006). Indeed, there is evidence for some weak purifying selection acting on synonymous sites in angiosperm mitogenomes (SLOAN and TAYLOR 2010; WYNN and CHRISTENSEN 2015). Selection on multinucleotide mutations may also affect observed synonymous substitution rates. There is a growing appreciation that clustered substitutions at adjacent sites can occur in a single mutational event (SCHRIDER *et al*. 2011; HARRIS and NIELSEN 2014) and that they can affect inferences of selection (VENKAT *et al*. 2018). It is very likely that some of the SNPs observed at adjacent sites in our analysis (Table S1) do not represent independent events. When such events occur in protein-coding genes, synonymous mutations may be removed by selection because they are linked to harmful mutations at adjacent nonsynonymous sites, whereas multinucleotide mutations in intergenic regions may remain relatively neutral. There are also mechanisms that may inflate substitution rate estimates in intergenic regions. For example, these regions often contain short, non-identical repeats that can undergo rare recombination events and create rearrangements (ARRIETA-MONTIEL *et al*. 2009; GUALBERTO and NEWTON 2017). Such recombination events can give the false impression that conventional nucleotide substitutions occurred because they create chimeric versions of similar but non-identical sequences.

Regardless of the causes of the small observed gap between substitution rates at synonymous sites vs. intergenic regions, it is clear that the magnitude of this difference is trivial relative to the wildly different rates of overall divergence observed between genes and the rest of the mitogenome in angiosperms. Indeed, it may simply reflect sampling variance as the small difference between intergenic regions and synonymous sites (0.0034 vs. 0.0027) is not even statistically significant (χ^2^ = 0.8; *P* = 0.37). While it is possible that certain mutational mechanisms preferentially act in intergenic regions and make them mutation ‘hotspots’, we favor an explanation based on strong selection on gene function, with region-specific mutation rates playing, at best, a secondary role in *A. thaliana* mitogenomes.

### Uneven copy number across angiosperm mitogenomes and implications for models of genome structure

Our analysis of a published sequencing dataset from MA lines (JIANG *et al*. 2014) and newly sequenced samples of purified mtDNA found evidence that coverage across the mitogenome is not constant and that it can change rapidly on generational timescales. Patterns of coverage variation were largely continuous (Figures 1 and 3), which contrasts with other commonly studied forms of copy number variation, in which germ-line segmental duplications or losses result in discrete shifts in coverage for specific regions of the genome (CONRAD *et al*. 2010). Our findings are relevant to previous work in the mitogenome of *Mimulus guttatus*, in which alternative recombination-mediated conformations showed evidence of heterogenous coverage, even in some cases where they were predicted to be part of the same subgenomic molecules (MOWER *et al*. 2012a). In addition, it has been shown that, disruption of specific nuclear genes involved in mitogenome replication, recombination, and repair can lead to preferential amplification or loss of certain genomic regions (SHEDGE *et al*. 2007; WALLET *et al*. 2015), and recent evidence indicates that mitogenome copy number can change in gene-specific ways across development in *Cucumis melo* (SHEN et al. 2019).

Other analyses of intraspecific mitogenome variation in systems such as *A. thaliana* (DAVILA et al. 2011), *Beta vulgaris* (DARRACQ *et al*. 2011), and *Zea mays* (ALLEN *et al*. 2007; DARRACQ *et al*. 2010) have generally focused on structural rearrangements resulting from repeat-mediated recombination. Indeed, at an even finer level, angiosperm mitogenomes are really a population of alternative structures that interconvert via recombination and coexist within cells and tissues in a single individual (PALMER and SHIELDS 1984; GUALBERTO and NEWTON 2017; KOZIK *et al*. 2019). As such, these structural rearrangements are arguably the most dynamic element of plant mtDNA evolution, and rapid shifts in the predominant structure (referred to as substoichiometric shifting) are often observed on very short generational timescales (ABDELNOOR *et al*. 2003; ARRIETA-MONTIEL and MACKENZIE 2011). However, when it comes to the MA-line analysis in this study, it is notable that it was copy number variation and not structural rearrangements for which we could detect significant divergence among lines. Therefore, in this case, it appears that copy number variation is the most rapidly diverging feature of the *A. thaliana* mitogenome, even though the general pattern of coverage is quite similar across lines (Figure 1) and there is known to be a persistent level of recombinational activity always going on beneath the surface. This interpretation is further supported by the fact that samples of the *A. thaliana* Col-0 accession from two different labs showed different patterns of copy number variation (Figures 1, 3, and S4). The divergence in copy number among lines did not appear to be entirely random, as we detected significant differences associated with salt-stress treatments, suggesting that the historical environment experienced in recent generations can have an effect in shaping the mitogenome landscape.

Angiosperm mitogenome sequencing projects typically report genome assemblies represented as a single circular structure, but it is widely accepted that this is an oversimplification resulting from mapping and that the physical form of angiosperm mtDNA involves complex branching structures (BENDICH 1993; SLOAN 2013; KOZIK *et al*. 2019). These branching structures likely reflect the activity of DNA replication, which is thought to be initiated by recombination-dependent mechanisms and not depend on a single origin of replication (CUPP and NIELSEN 2014). In addition to findings from more direct observations of the physical form of mtDNA molecules (BENDICH 1996; BACKERT and BORNER 2000), coverage patterns in previous sequencing efforts have been interpreted as evidence against a ‘master circle’ as the predominant form of the mitogenome (MOWER *et al*. 2012a).

By itself, copy number variation is not definitive evidence against a simple circular organization in *A. thaliana*. Bacterial genomes are circular structures but can still exhibit quantitative variation in coverage across the genome when DNA is sampled from actively dividing cultures, with copy number decreasing from the origin of replication to the terminus of replication. Indeed, analyzing sequencing coverage of bacterial genomes can be an effective way to identify the location of the origin of replication and measure the replication rate of bacteria (BROWN *et al*. 2016). Nevertheless, we contend that the combination of heterogeneous coverage and evidence for rapid shifts in copy number variation is unlikely to be explained by a simple circular model with preferential amplification at origin(s) of replication within the circle, especially when viewed in the light of existing evidence against the master circle as a predominant genome form. Instead, our results suggest that the complex physical structure of angiosperm mitogenomes creates opportunities for differential amplification of subgenomic regions in a dynamic way that does not occur in simpler mitogenomes like those found in bilaterian animals. In addition to the rapid and large changes in the frequencies of mitogenome structural conformations associated with substoichiometric shifting, angiosperms appear to be subject to more pervasive low-level fluctuations in copy numbers of local regions within the genome.

## ACKNOWLEDGEMENTS

We thank Jeff Mower for helpful discussion and motivation to use intraspecific variation to analyze contrasting patterns of evolution between genic and intergenic regions in plant mitogenomes. This work was supported by a grant from the NIH (R01 GM118046) and an NSF NRT GAUSSI Graduate Fellowship (DGE-1450032).

---

**Figure S1.**
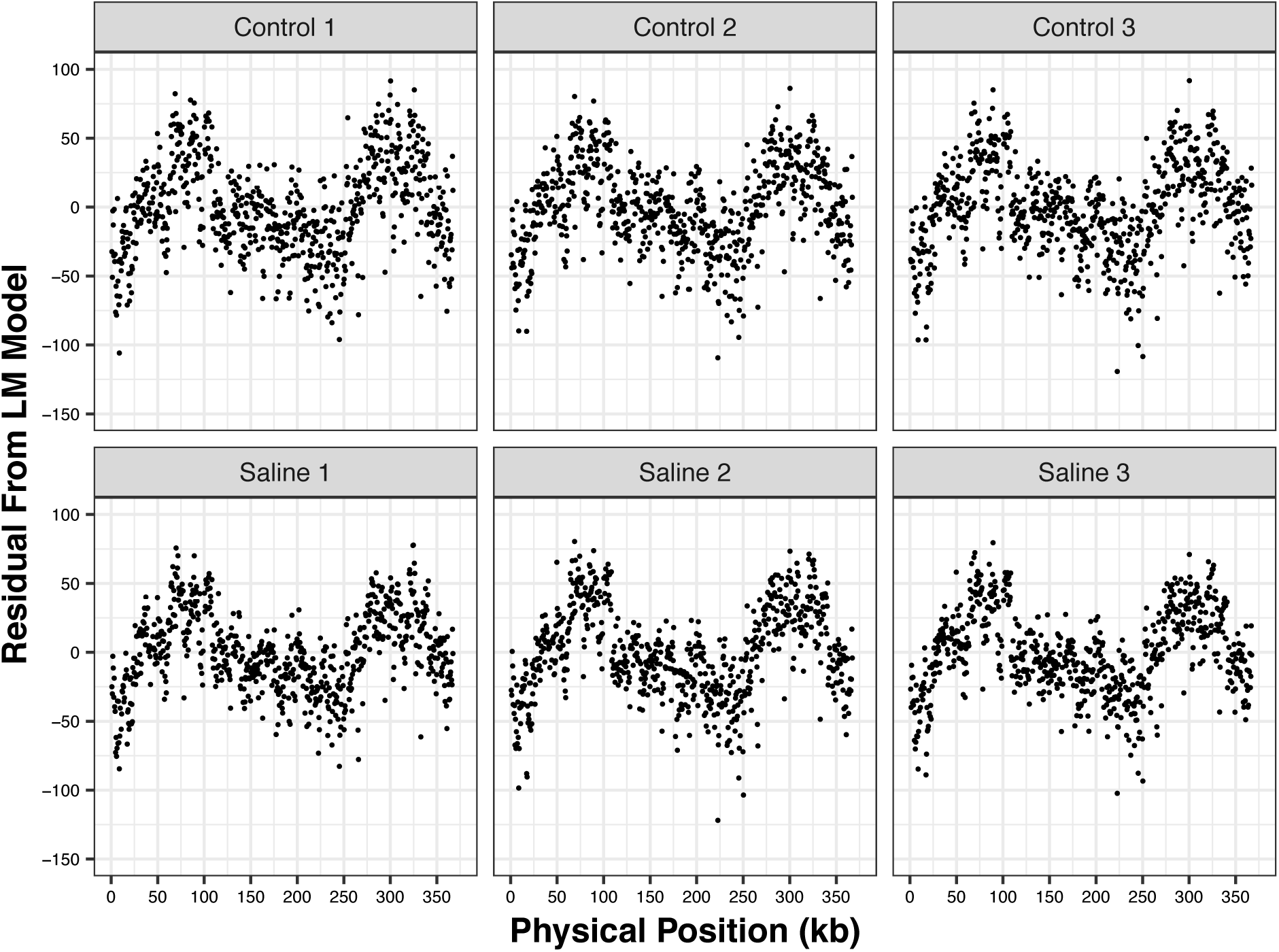
Sequencing coverage variation across mitogenome of *Arabidopsis thaliana* mutation accumulation lines as measured by the residuals from a model accounting for sequencing bias due to nucleotide composition. Each panel represents an average of three biological replicates.

**Figure S2.**
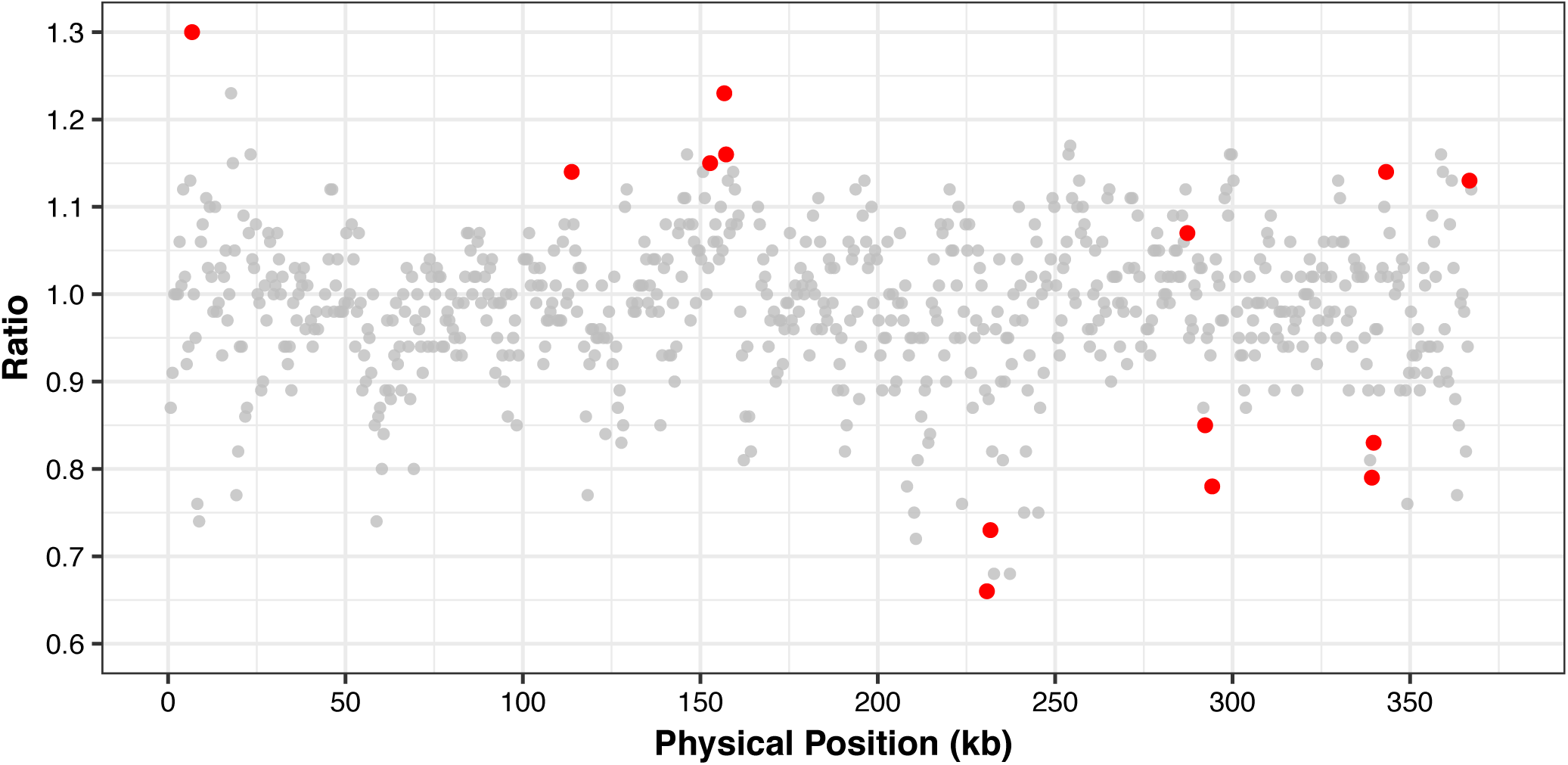
Divergence in region-specific mitogenome copy number in salt-stressed vs. control mutation accumulation lines. Values are expressed as a ratio of values for all salt-stressed and all control lines. These are the same data depicted in Figure 2 except that value were calculated as the genome-wide mean CPMM plus the residual from a linear model that accounts for sequencing bias due to nucleotide composition. Windows that deviate significantly from a ratio of 1 after false-discovery-rate correction are highlighted in red. CPMM: counts per million mapped reads.

**Figure S3.**
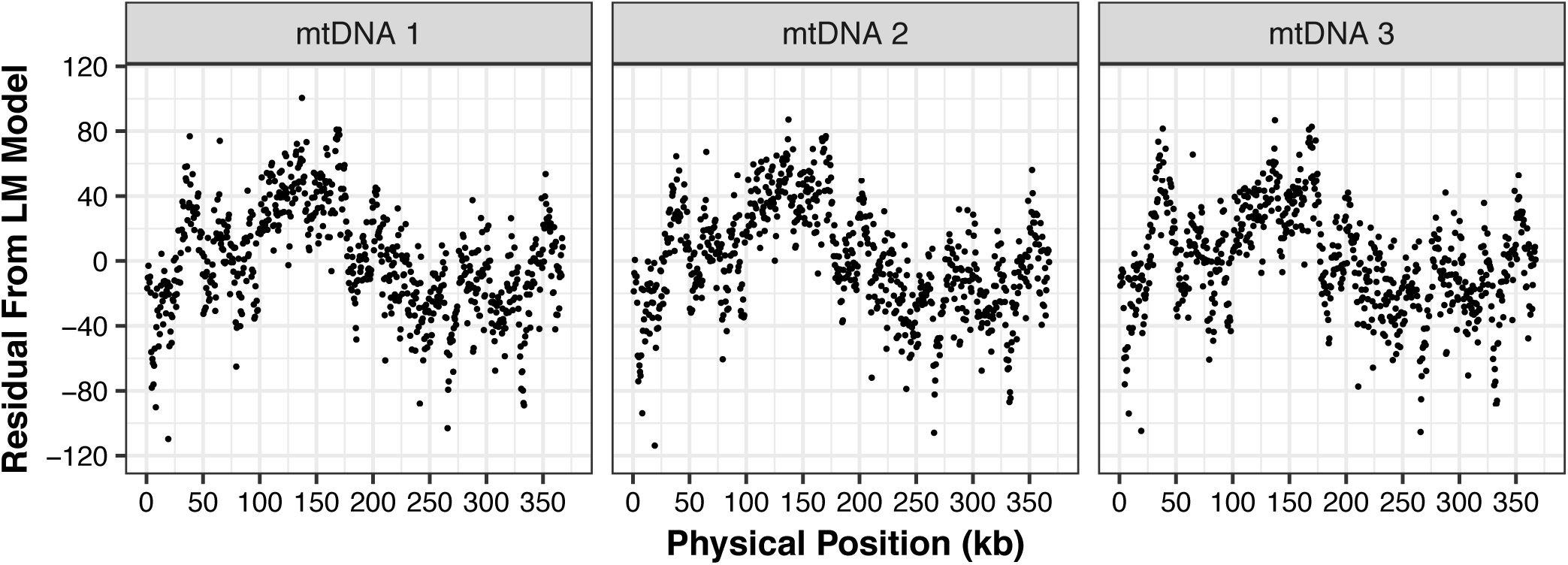
Sequencing coverage variation across the mitogenome for three purified mtDNA samples from *Arabidopsis thaliana* as measured by the residuals from a model accounting for sequencing bias due to nucleotide composition.

**Figure S4.**
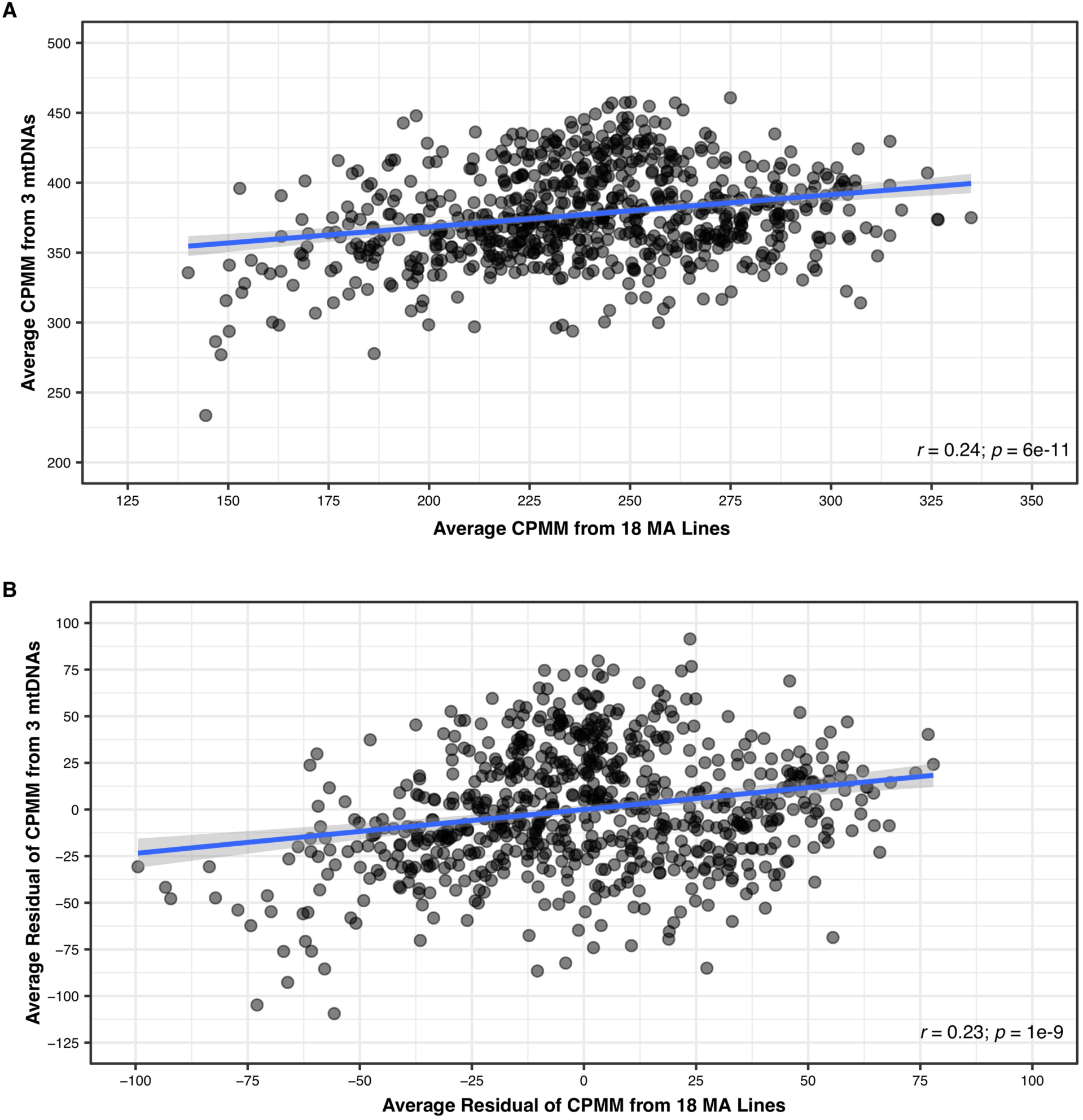
Correlation between mitogenome coverage from published Col-0 mutation accumulation lines (JIANG *et al*. 2014) and purified mtDNA from our Col-0 lab line. Each point represents a 500-bp window. Coverage is expressed as either (A) raw copies per million mapped reads (CPMM) or (B) residuals from a model that accounts for sequencing bias due to nucleotide composition.

## REFERENCES

Abdelnoor, R. V., R. Yule, A. Elo, A. C. Christensen, G. Meyer-Gauen et al., 2003 Substoichiometric shifting in the plant mitochondrial genome is influenced by a gene homologous to MutS. Proceedings of the National Academy of Sciences 100: 5968–5973.

Aird, D., M. G. Ross, W. S. Chen, M. Danielsson, T. Fennell et al., 2011 Analyzing and minimizing PCR amplification bias in Illumina sequencing libraries. Genome biology 12: R18.

Allen, J. O., C. M. Fauron, P. Minx, L. Roark, S. Oddiraju et al., 2007 Comparisons among two fertile and three male-sterile mitochondrial genomes of maize. Genetics 177: 1173–1192.

Alonso-Blanco, C., J. Andrade, C. Becker, F. Bemm, J. Bergelson et al., 2016 1,135 genomes reveal the global pattern of polymorphism in *Arabidopsis thaliana*. Cell 166: 481–491.

Arrieta-Montiel, M. P., and S. A. Mackenzie, 2011 Plant mitochondrial genomes and recombination, pp. 65–82 in Plant Mitochondria, edited by F. Kempken. Springer Verlag, New York.

Arrieta-Montiel, M. P., V. Shedge, J. Davila, A. C. Christensen and S. A. Mackenzie, 2009 Diversity of the Arabidopsis mitochondrial genome occurs via nuclear-controlled recombination activity. Genetics 183: 1261–1268.

Backert, S., and T. Borner, 2000 Phage T4-like intermediates of DNA replication and recombination in the mitochondria of the higher plant Chenopodium album (L.). Current genetics 37: 304–314.

Bendich, A. J., 1993 Reaching for the ring: the study of mitochondrial genome structure. Current genetics 24: 279–290.

Bendich, A. J., 1996 Structural analysis of mitochondrial DNA molecules from fungi and plants using moving pictures and pulsed-field gel electrophoresis. Journal of Molecular Biology 255: 564–588.

Benjamini, Y., and Y. Hochberg, 1995 Controlling the False Discovery Rate: A Practical and Powerful Approach to Multiple Testing. Journal of the Royal Statistical Society.Series B (Methodological) 57: 289–300.

Brown, C. T., M. R. Olm, B. C. Thomas and J. F. Banfield, 2016 Measurement of bacterial replication rates in microbial communities. Nature Biotechnology 34: 1256–1263.

Brown, W. M., M. George and A. C. Wilson, 1979 Rapid evolution of animal mitochondrial DNA. Proceedings of the National Academy of Sciences 76: 1967–1971.

Chamary, J. V., J. L. Parmley and L. D. Hurst, 2006 Hearing silence: non-neutral evolution at synonymous sites in mammals. Nature reviews.Genetics 7: 98–108.

Christensen, A. C., 2013 Plant mitochondrial genome evolution can be explained by DNA repair mechanisms. Genome Biology and Evolution 5: 1079–1086.

Christensen, A. C., 2014 Genes and junk in plant mitochondria-repair mechanisms and selection. Genome Biology and Evolution 6: 1448–1453.

Conrad, D. F., D. Pinto, R. Redon, L. Feuk, O. Gokcumen et al., 2010 Origins and functional impact of copy number variation in the human genome. Nature 464: 704–712.

Cupp, J. D., and B. L. Nielsen, 2014 DNA replication in plant mitochondria. Mitochondrion 19: 231–237.

Darracq, A., J. S. Varre, L. Marechal-Drouard, A. Courseaux, V. Castric et al., 2011 Structural and content diversity of mitochondrial genome in beet: a comparative genomic analysis. Genome biology and evolution 3: 723–736.

Darracq, A., J. S. Varre and P. Touzet, 2010 A scenario of mitochondrial genome evolution in maize based on rearrangement events. BMC genomics 11: 233.

Davila, J. I., M. P. Arrieta-Montiel, Y. Wamboldt, J. Cao, J. Hagmann et al., 2011 Double-strand break repair processes drive evolution of the mitochondrial genome in Arabidopsis. BMC biology 9: 64.

Doyle, J. J., and J. L. Doyle, 1987 A rapid DNA isolation procedure for small quantities of fresh leaf tissue. Phytochemical bulletin 19: 11–15.

Drouin, G., H. Daoud and J. Xia, 2008 Relative rates of synonymous substitutions in the mitochondrial, chloroplast and nuclear genomes of seed plants. Molecular Phylogenetics and Evolution 49: 827–831.

Ellis, J., 1982 Promiscuous DNA--chloroplast genes inside plant mitochondria. Nature 299: 678–679.

Goremykin, V. V., P. J. Lockhart, R. Viola and R. Velasco, 2012 The mitochondrial genome of Malus domestica and the import-driven hypothesis of mitochondrial genome expansion in seed plants. The Plant Journal 71: 615–626.

Gualberto, J. M., and K. J. Newton, 2017 Plant mitochondrial genomes: dynamics and mechanisms of mutation. Annual Review of Plant Biology 68: 225–252.

Halligan, D. L., and P. D. Keightley, 2009 Spontaneous mutation accumulation studies in evolutionary genetics. Annual Review of Ecology, Evolution, and Systematics 40: 151–172.

Hanawalt, P. C., and G. Spivak, 2008 Transcription-coupled DNA repair: two decades of progress and surprises. Nature Reviews Molecular Cell Biology 9: 958–970.

Harris, K., and R. Nielsen, 2014 Error-prone polymerase activity causes multinucleotide mutations in humans. Genome Research 24: 1445–1454.

Hazkani-Covo, E., R. M. Zeller and W. Martin, 2010 Molecular poltergeists: mitochondrial DNA copies (numts) in sequenced nuclear genomes. PLoS Genetics 6: e1000834.

Jiang, C., A. Mithani, E. J. Belfield, R. Mott, L. D. Hurst et al., 2014 Environmentally responsive genome-wide accumulation of de novo Arabidopsis thaliana mutations and epimutations. Genome Research 24: 1821–1829.

Kozik, A., B. Rowan, D. Lavelle, L. Berke, M. E. Schranz et al., 2019 The alternative reality of plant mitochondrial DNA. bioRxiv: 564278.

Kubo, T., and K. J. Newton, 2008 Angiosperm mitochondrial genomes and mutations. Mitochondrion 8: 5–14.

Langmead, B., and S. L. Salzberg, 2012 Fast gapped-read alignment with Bowtie 2. Nature Methods 9: 357–359.

Li, H., B. Handsaker, A. Wysoker, T. Fennell, J. Ruan et al., 2009 The Sequence Alignment/Map format and SAMtools. Bioinformatics (Oxford, England) 25: 2078–2079.

Lonsdale, D. M., T. Brears, T. P. Hodge, S. E. Melville and W. H. Rottmann, 1988 The plant mitochondrial genome: homologous recombination as a mechanism for generating heterogeneity. Philosophical Transactions of the Royal Society of London.B, Biological Sciences 319: 149–163.

Martin, M., 2011 Cutadapt removes adapter sequences from high-throughput sequencing reads. EMBnet.journal 17: 10–12.

McKenna, A., M. Hanna, E. Banks, A. Sivachenko, K. Cibulskis et al., 2010 The Genome Analysis Toolkit: a MapReduce framework for analyzing next-generation DNA sequencing data. Genome Research 20: 1297–1303.

Mower, J. P., A. L. Case, E. R. Floro and J. H. Willis, 2012a Evidence against equimolarity of large repeat arrangements and a predominant master circle structure of the mitochondrial genome from a monkeyflower (Mimulus guttatus) lineage with cryptic CMS. Genome Biology and Evolution 4: 670–686.

Mower, J. P., D. B. Sloan and A. J. Alverson, 2012b Plant mitochondrial diversity – the genomics revolution, pp. 123–144 in Plant Genome Diversity, edited by J. F. Wendel. Springer, Vienna.

Nielsen, R., 2005 Molecular signatures of natural selection. Annual Review of Genetics 39: 197–218.

Palmer, J. D., and C. R. Shields, 1984 Tripartite structure of the Brassica campestris mitochondrial genome. Nature 307: 437–440.

Rice, D. W., A. J. Alverson, A. O. Richardson, G. J. Young, M. V. Sanchez-Puerta et al., 2013 Horizontal transfer of entire genomes via mitochondrial fusion in the angiosperm Amborella. Science 342: 1468–1473.

Robinson, J. T., H. Thorvaldsdóttir, A. M. Wenger, A. Zehir and J. P. Mesirov, 2017 Variant review with the integrative genomics viewer. Cancer Research 77: e31–e34.

Schrider, D. R., J. N. Hourmozdi and M. W. Hahn, 2011 Pervasive multinucleotide mutational events in eukaryotes. Current Biology 21: 1051–1054.

Shedge, V., M. Arrieta-Montiel, A. C. Christensen and S. A. Mackenzie, 2007 Plant mitochondrial recombination surveillance requires unusual RecA and MutS homologs. The Plant Cell 19: 1251–1264.

Shen, J., Y. Zhang, M. J. Havey and W. Shou, 2019 Copy numbers of mitochondrial genes change during melon leaf development and are lower than the numbers of mitochondria. Horticulture Research 6: 95.

Sloan, D. B., 2013 One ring to rule them all? Genome sequencing provides new insights into the ‘master circle’ model of plant mitochondrial DNA structure. New Phytologist 200: 978–985.

Sloan, D. B., J. C. Havird and J. Sharbrough, 2017 The on-again, off-again relationship between mitochondrial genomes and species boundaries. Molecular Ecology 26: 2212–2236.

Sloan, D. B., K. Muller, D. E. McCauley, D. R. Taylor and H. Storchova, 2012 Intraspecific variation in mitochondrial genome sequence, structure, and gene content in Silene vulgaris, an angiosperm with pervasive cytoplasmic male sterility. The New Phytologist 196: 1228–1239.

Sloan, D. B., and D. R. Taylor, 2010 Testing for selection on synonymous sites in plant mitochondrial DNA: the role of codon bias and RNA editing. Journal of Molecular Evolution 70: 479–491.

Sloan, D. B., and Z. Wu, 2014 History of plastid DNA insertions reveals weak deletion and at mutation biases in angiosperm mitochondrial genomes. Genome Biology and Evolution 6: 3210–3221.

Sloan, D. B., Z. Wu and J. Sharbrough, 2018 Correction of persistent errors in Arabidopsis reference mitochondrial genomes. Plant Cell 30: 525–527.

Small, I. D., P. G. Isaac and C. J. Leaver, 1987 Stoichiometric differences in DNA molecules containing the atpA gene suggest mechanisms for the generation of mitochondrial genome diversity in maize. The EMBO journal 6: 865–869.

Stajich, J. E., D. Block, K. Boulez, S. E. Brenner, S. A. Chervitz et al., 2002 The Bioperl toolkit: Perl modules for the life sciences. Genome research 12: 1611–1618.

Stupar, R. M., J. W. Lilly, C. D. Town, Z. Cheng, S. Kaul et al., 2001 Complex mtDNA constitutes an approximate 620-kb insertion on Arabidopsis thaliana chromosome 2: implication of potential sequencing errors caused by large-unit repeats. Proceedings of the National Academy of Sciences of the United States of America 98: 5099–5103.

van Dijk, E. L., Y. Jaszczyszyn and C. Thermes, 2014 Library preparation methods for next-generation sequencing: tone down the bias. Experimental Cell Research 322: 12–20.

Venkat, A., M. W. Hahn and J. W. Thornton, 2018 Multinucleotide mutations cause false inferences of lineage-specific positive selection. Nature Ecology & Evolution 2: 1280–1288.

Wallet, C., M. Le Ret, M. Bergdoll, M. Bichara, A. Dietrich et al., 2015 The RECG1 DNA translocase is a key factor in recombination surveillance, repair, and segregation of the mitochondrial DNA in Arabidopsis. Plant Cell 27: 2907–2925.

Warren, J. M., M. P. Simmons, Z. Wu and D. B. Sloan, 2016 Linear plasmids and the rate of sequence evolution in plant mitochondrial genomes. Genome Biology and Evolution 8: 364–374.

Wolfe, K. H., W. H. Li and P. M. Sharp, 1987 Rates of nucleotide substitution vary greatly among plant mitochondrial, chloroplast, and nuclear DNAs. Proceedings of the National Academy of Sciences of the United States of America 84: 9054–9058.

Wynn, E. L., and A. C. Christensen, 2015 Are synonymous substitutions in flowering plant mitochondria neutral? Journal of Molecular Evolution 81: 131–135.

Yang, Z., and A. D. Yoder, 1999 Estimation of the transition/transversion rate bias and species sampling. Journal of Molecular Evolution 48: 274–283.

Zhu, A., W. Guo, K. Jain and J. P. Mower, 2014 Unprecedented heterogeneity in the synonymous substitution rate within a plant genome. Molecular Biology and Evolution 31: 1228–1236.

